# Enteric neural crest development in *Astyanax mexicanus* surface fish and cavefish

**DOI:** 10.1101/2025.02.24.639940

**Authors:** Pavani Ponnimbaduge Perera, Kaitlyn Webster, Misty R Riddle

**Affiliations:** University of Nevada, Reno, Reno, NV, USA; Harvard Medical School, Boston, MA, USA

**Author notes:** Corresponding author Misty Riddle University of Nevada, Reno 1664 N Virginia St Reno, NV 89557.

**Keywords:** Enteric nervous system, *Astyanax mexicanus*, enteric neural crest, evolution, development

## Abstract

The enteric nervous system (ENS) regulates gastrointestinal (GI) functions such as secretion, blood flow, and motility, yet how its structure and function evolve with dietary adaptations remains unclear. *Astyanax mexicanus*, a teleost fish with surface and cave morphotypes, offers a model to explore these changes; cavefish exhibit altered GI motility and transit that may help them adapt to their unique diet. We compared early ENS development in surface fish and cavefish, tracking enteric neural crest cell (ENCC) migration and differentiation using *phox2bb* and HuC/D markers. We found that ENCCs reach the gut by 36 hours post-fertilization (hpf) in both morphotypes but migrate and differentiate more rapidly along the gut tube in cavefish. To explore the genetic basis of this difference, we used available genomic datasets to compare the predicted peptide sequences of genes important for ENS development in other species and identified mutations that could impact protein function, for example in the endothelin signaling genes important for ENCC migration and differentiation. We specifically examined the expression of *endothelin-3 (edn3)* and *endothelin receptor-b (ednrb)* during ENCC migration and found that the localization of *edn3*, but not *ednrb*, is consistent with a role in ENS development. Overall, our findings position *A. mexicanus* as a model for studying evolution of ENS development.

## Introduction

Neural crest cells (NCCs) are multipotent migratory cells that originate from the neural tube during vertebrate embryonic development and give rise to a diverse array of cell types that contribute to the cranium, peripheral nervous system, integument, and gastrointestinal tract (GI) (reviewed in^1–3^). Diversification of neural crest cell development has contributed to vertebrate evolution by permitting formation of novel cell types and structures^4^. The enteric nervous system (ENS), an autonomous network of interconnected neurons and glia embedded in the GI tract, develops primarily from NCCs^5–8^. The ENS regulates motility, blood flow, and secretion of hormones in the gut^9–11^ and is essential for proper digestion of food and elimination of waste. Vertebrates display dramatic differences in GI tract development and digestive physiology that are important for maximizing energy extraction from distinct diets^12^. However, little is known about how the vertebrate ENS evolves as animals adapt to new environments. Investigating natural variation in ENS development and how it impacts gut physiology is challenging as collecting embryos from diverse vertebrates for comparative studies may not be feasible.

The Mexican tetra, *Astyanax mexicanus*, is a promising model system to study vertebrate ENS evolution and development. *A. mexicanus* is a species of teleost fish that exists as a river population (Surface fish, Fig 1A, C) and eyeless cave populations (Cavefish, Fig 1B, D) named for the caves they inhabit in the Sierra de El Abra region of Northeastern Mexico (i.e., Pachón, Molino, Tinaja). The river and cave populations evolved with dramatically different resources; the caves are perpetually dark making cavefish dependent on food brought in by bats or seasonal floods compared to surface fish that have access to plants and insects^13^. Cavefish populations that evolved independently from surface fish have converged on similar morphology, behavior, and metabolism permitting investigation of parallel evolution (reviewed in^14,15^). The populations are easy to maintain and breed in the laboratory producing hundreds of embryos with each spawning event that can be used to compare embryonic development. The cave and surface morphotypes are also interfertile which allows researchers to genetically map morphological, physiological, and behavior traits^16–21^. Furthermore, the genomes are sequenced and annotated^22,23^ and methods for transgenesis and gene editing are commonly used^24,25^ allowing detailed investigations into development and gene function.

**Figure 1.**
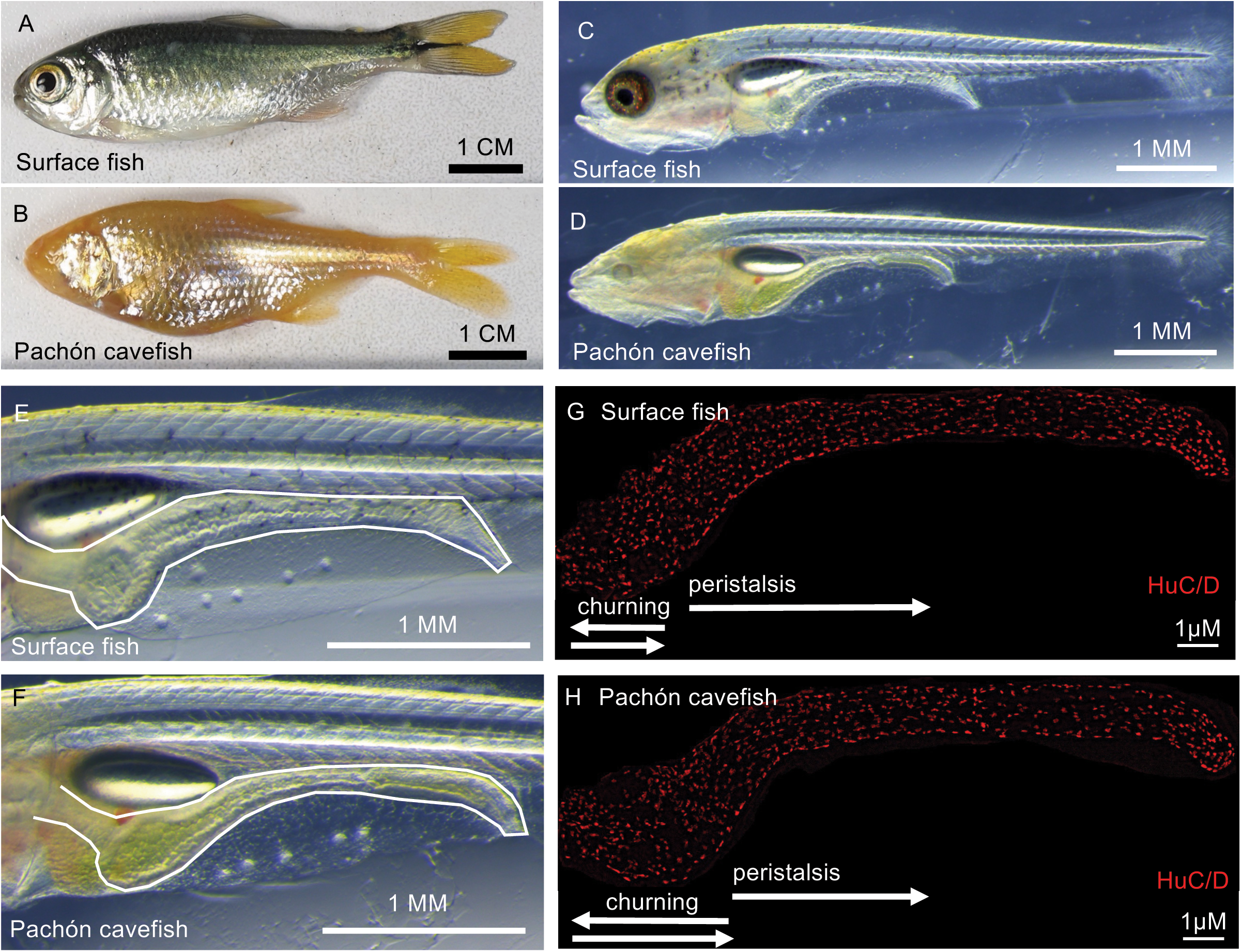
**Astyanax mexicanus as a model to study enteric nervous system (ENS) development and evolution**. The Mexican tetra, *A. mexicanus,* is a single species of fish that exists as river-adapted surface fish (A) and cave-adapted cavefish named for the caves they inhabit (e.g. Pachón cavefish, B). Post-larval surface fish (C) and cavefish (D) are transparent, permitting visualization of the gastrointestinal tract (white outline, E, F) within intact animals. Immunostaining using a pan-neuronal antibody reveals the entire ENS in 12.5-day-old surface fish (G) and Pachón cavefish guts. Surface fish and Pachón cavefish display differences in gastrointestinal motility at this stage with Pachón having a greater portion of the gut that produces waves traveling retrograde and anterograde directions (white arrows, churning) and surface fish having a greater portion of the gut that produces only anterograde waves (white arrow, peristalsis). At this stage, surface fish and cavefish do not display differences in total enteric neuron number (∼700 neurons).

Previous research comparing the gut of post-larval *A. mexicanus* surface fish and Pachón cavefish (Fig 1C-F) revealed differences in GI motility^26^. Pachón cavefish exhibit broader regions of churning motility and slower digesta transit from the stomach-like region to the midgut^26^. This delayed transit may help cavefish extract nutrients in resource-poor caves. At the stage when motility differences appear, the ENS comprises a single neuronal layer (∼700 neurons) sandwiched between two muscle layers, with no differences in total neuron number between morphotypes (Fig 1G,H)^26^. Comparing ENS development could uncover the basis for altered motility and advance our understanding of vertebrate ENS evolution.

Here we compared early ENS development between surface fish and Pachón cavefish. First, we constructed a developmental timeline of enteric neural crest cell (ENCCs) migration and differentiation using marker genes *phox2bb* and HuC/D, which label ENCCs and differentiated enteric neurons, respectively. Our results indicate that ENCCs migrate to the *A. mexicanus* gut by 36 hours post-fertilization (hpf), and completely colonize the gut by 60 hpf in both surface fish and Pachón cavefish. However, migration and differentiation along the gut tube may occur more slowly in surface fish; at 48hpf, we observed that *phox2bb*-expressing cells had dispersed closer to the end of the gut tube in Pachón cavefish compared to surface fish, and that only Pachón cavefish have HuC/D-expressing cells at this stage. Second, we examined the expression of genes essential for normal ENCC migration and differentiation in other species, *endothelin-3* (*edn3*) and *endothelin receptor b* (*ednrb*). We found that *edn3* is expressed in *A. mexicanus* in cells ventral to the developing gut tube, ahead of the migratory ENCCs, consistent with a role in facilitating ENCC migration. However, we did not find cells that express both *phox2b* and *ednrb*. Third, we compared the coding sequences of known regulators of ENS development between surface fish and Pachón cavefish and identified mutations that may impact protein functions. Our findings shed light on the potential origins of motility differences between surface fish and cavefish, and position *A. mexicanus* to investigate ENS development and evolution.

## Materials and Methods

### Fish husbandry

Laboratory raised adult Surface fish and Pachón cavefish were bred using natural spawning. The embryos used for hybridization chain reaction were collected at the single cell stage directly after spawning, incubated at 25°C, and fixed in 4% PFA at 24hpf, 27hpf, 36 hpf, 48 hpf and 60 hpf. The samples were dehydrated and stored in methanol. For samples collected between 3-3.5dpf, the time of spawning was estimated using a developmental staging table ^27^ and used to determine the larvae age at the time of collection.

### Immunohistochemistry

Immunohistochemistry was done in *A. mexicanus* larvae according to the protocol described^28^. The primary antibodies used are 1:250 and 1:500 HuC/D Neuronal Protein Mouse Monoclonal Antibody (Invitrogen, A-21271). The secondary antibodies used are 1:500 Goat Anti-Mouse IgG H&L (Alexa Fluor® 488) preadsorbed (ab150117) and 1:1000 Goat Anti-Mouse IgG H&L (Alexa Fluor® 647) preadsorbed (ab150119).

### Hybridization chain reaction

HCR probe sets and amplifiers were ordered from Molecular Instruments Inc. HCR probes with B isoforms include *A. mexicanus sox10*-B1 (ENSAMXG00000036184)*, phox2bb*-B4 (ENSAMXG00000043803), *ednrba*-B2 (ENSAMXG00000036587), *edn3*-B3 (ENSAMXG00000032596). HCR was done in *A. mexicanus* larvae according to the protocol described^29^.

### Microscopy

Leica Stellaris 5 confocal Microscope, Keyence BZ-9000 fluorescence microscope, Leica THUNDER 3D Tissue Upright Microscope, and Leica M165SC stereomicroscope were used for imaging fish embryos and larvae.

### Comparative genomics

We used the annotated surface fish genome (AstMex3_surface, NCBI Assembly GCA_023375975.1, May 2022) and annotated Pachón cavefish genome (AMEX_1.1, NCBI Assembly GCA_019721115.1, Aug 2021) to compare the predicted protein sequences of genes involved in ENS development in zebrafish. For each gene, we downloaded the predicted protein sequence of the closest zebrafish ortholog, using the longest splice isoform for alignments. We used SnapGene software with Needle-man Wunsch global alignment method for pairwise alignments, and MUSCLE (Multiple Sequence Comparison by Log-Expectation) algorithm for multiple alignments. Sequence changes are annotated relative to the surface fish peptide.

## Results and discussion

### ENS developmental timeline reveals evidence of accelerated ENCC migration in Pachón cavefish

The enteric neural crest cells (ENCCs) in zebrafish originate from the vagal neural crest, located near somites 1–7 ^30^. These cells migrate ventrally, guided by interactions with surrounding tissues, and enter the gut via the anterior region, where they proliferate, migrate rostrocaudally along the gut mesenchyme, and differentiate into neurons and glia to form the enteric nervous system (ENS) ^31–33^. In zebrafish, ENCCs reach the foregut by 36hpf and complete the migration to the end of the gut by 72hpf^33–35^. They differentiate in a wave from anterior to posterior with enteric neurons observable in the foregut as early as 54hpf^32,33^.

We utilized the zebrafish ENCC marker genes *sox10* and *phox2b* to compare ENCC migration between *A. mexicanus* surface fish and Pachón cavefish. In zebrafish, migratory vagal neural crest cells express *sox10* and *phox2b,* while expression of only *phox2b* persists in ENCCs that are fated to become enteric neurons^36^. The closest orthologs to these genes in *A. mexicanu*s are *sox10* (ENSAMXG00000036184) and *phox2bb* (ENSAMXG00000043803). In both *A. mexicanus* morphotypes, we observed *sox10* and *phox2bb* transcripts in cells posterior to the otic vesicle at 24-27hpf, and at 36hpf, *phox2bb* transcripts in cells within the foregut region (Fig 2A-B). Combined, these results suggest that migration of vagal neural crest cells to the foregut occurs between 24-36hpf in both surface fish and cavefish. In contrast, migration of ENCCs along the gut mesenchyme occurs at different rates (Fig 2L). At 48 hpf we observed *phox2bb*-positive cells only in the anterior portion of the gut in surface fish while in Pachón cavefish, *phox2bb*-positive cells were detected within the anterior and proximal gut (Fig. 2E-F). In 60 hpf larvae, we observed cells containing *phox2bb* transcripts throughout the entire gut in both morphotypes, suggesting ENCCs have reached the end of the hindgut in surface fish at this stage (Fig 2G-H). These findings are consistent with more rapid migration of ENCCs along the gut tube in Pachón cavefish.

**Figure 2.**
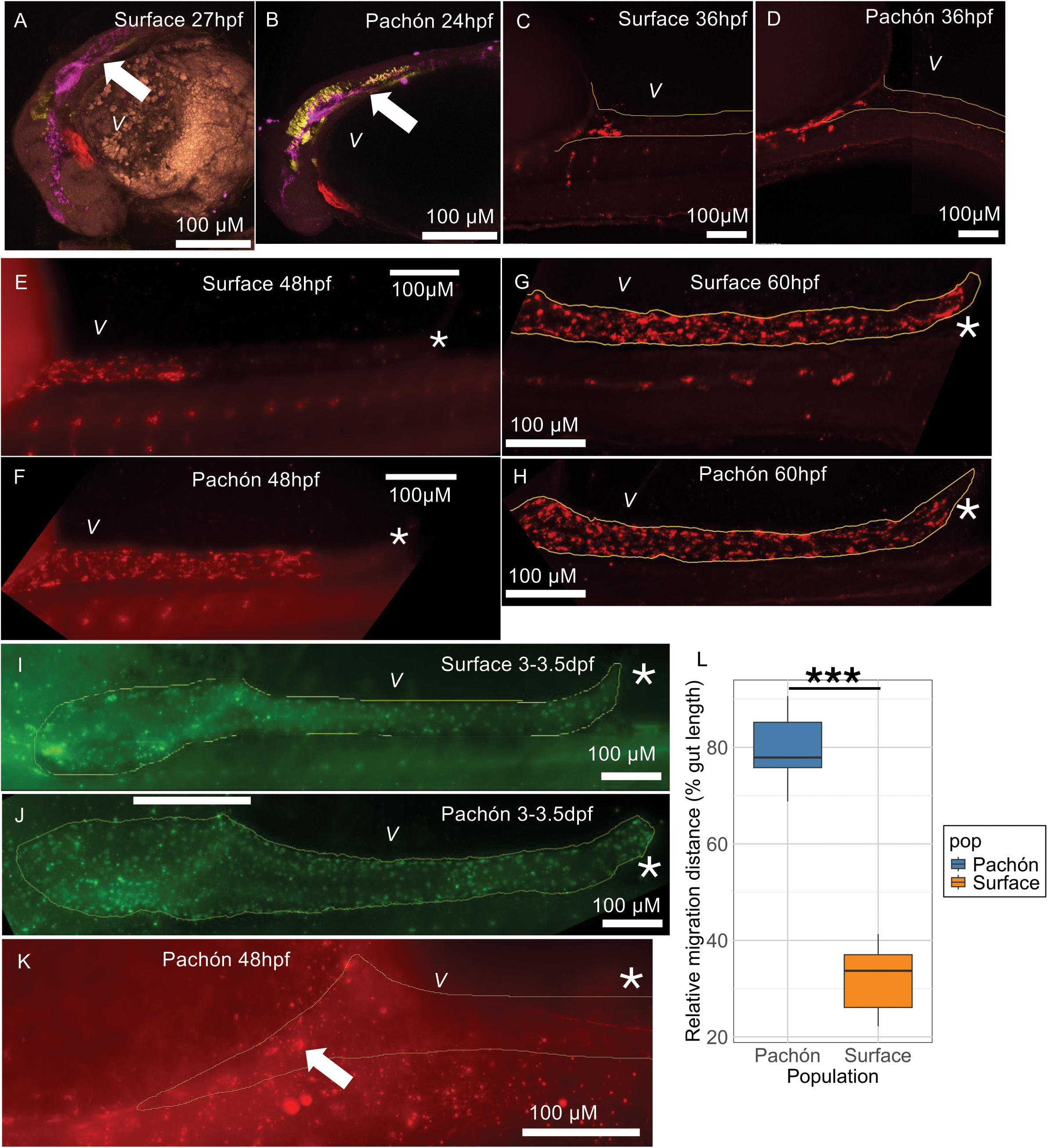
Comparison of enteric neural crest cell migration and differentiation between *Astyanax mexicanus* surface fish and cavefish using hybridization chain reaction and immunohistochemistry. Migratory vagal neural crest cell population (magenta – *sox10*, yellow – *phox2b,* white arrow) can be seen posterior to the otic vesicles at 24-27 surface fish (A) and Pachón cavefish (B). *phox2b*+ ENCCs are seen in the anterior gut of surface fish (C) and Pachón cavefish (D) at 36 hpf. *phox2b*+ ENCCs dispersed throughout the gut tube at 48 hpf in Pachón cavefish (F) but not surface fish (E). *phox2b+* ENCCs dispersed throughout the gut tube at 60 hpf in both surface fish (G) and Pachón cavefish (H). HuC/D+ cells are observed throughout the entire gut of surface fish (I) and Pachón cavefish at 72-84 hpf. (J) (L) Quantification of *phox2b+* ENCCs migratory distance as a percentage of gut length in surface fish and Pachón cavefish at 48 hpf (n=5) ***p<0.001, students two-tailed *t*-test. (K) At 48 hpf HuC/D+ cells are only seen in Pachón cavefish (white arrow). Yellow outline – GI tract borders. “*” Posterior lateral end of the larval fish gut. *“V”* Ventral side of the fish.

In addition to more rapid migration, we also found evidence of earlier differentiation of ENCCs in Pachón cavefish. We observed cells expressing the differentiated neuronal marker HuC/D in Pachón cavefish, but not surface fish at 48hpf (Fig 2K, n= 2 out of 4 Pachón samples versus 0 out of 4 surface fish samples). The cells were only seen in the anterior region of the gut consistent with the process of ENCC development in zebrafish, where migration and differentiation proceed in a wave from anterior to posterior ^33^. By 72-84hpf, differentiated neurons were observed throughout the entire gut in both morphotypes (Figure 2I-J).

Localized gut contractions can be observed at 5.5 dpf in both morphotypes when the larvae start actively feeding and have a functioning digestive system^26,27^. Coordinated motility patterns however are not observed until 8.5 dpf ^26^. At this stage, differences in motility between surface fish and cavefish are first apparent, with Pachón cavefish displaying a greater proportion of the gut that exhibits churning motility^26^ (Fig 1G,H). In zebrafish, genetic manipulation of multiple signaling pathways has been shown to delay ENCC migration, ultimately resulting in a reduction in enteric neurons in larvae and altered GI motility (reviewed in^37^). Total enteric neuron number is not significantly different between cavefish and surface fish despite differences in motility suggesting other factors, such as neuronal subtype diversity, could be important to consider. How early differences in ENCC migration and differentiation between morphotypes impacts ENS differentiation and gut motility is currently unclear. Future comparisons using cavefish as “natural mutants” may however lead to insights in this area and expand our understanding of ENS development.

### Comparison of endothelin signaling gene sequence and expression in A. mexicanus

Endothelin signaling is essential for proper ENCC migration and differentiation in humans, mice, and zebrafish^38–40^. During ENCC migration, the ligand Edn3 is secreted by the gut mesenchyme^41^. Migrating ENCCs express the Edn3 receptor, Ednrb, which is also expressed at low levels in gut mesenchymal cells^42,43^. Edn3 binding to the receptor initiates a signaling cascade^44^ that promotes ENCCs proliferation and migration while inhibiting differentiation^38,39,45^. Edn3-mediated inhibition of differentiation maintains the ENCCs in an uncommitted state^46^, thereby promoting the even colonization of NCC along the GI tract. Mice that carry mutations in the *edn3* and *ednrb* genes lack enteric neurons in the hindgut and develop megacolon^47–49^. In humans, mutations in EDN3 and EDNRB result in Hirschsprung’s disease, a congenital birth defect characterize by lack of enteric neurons in the distal bowel and bowel obstruction^50 51^.

Considering the importance of the endothelin signaling pathway in regulating ENCC migration, we next examined whether mutations or altered expression of *edn3* or *ednrb* may occur in Pachón cavefish which have accelerated migration. Surface fish have two *endothelin receptor B* genes: *ednrba* located on chromosome 21 that is orthologous to zebrafish *ednrba* (NCBI ID103031642, ENSDARG00000089334) and *ednrb* located on chromosome 11 that is orthologous to zebrafish *ednrbb (*NCBI ID103043104, ENSDARG00000053498). In Pachón cavefish, the Ednrba protein has a point mutation, changing leucine to phenylalanine (L238F) in the fourth extracellular loop, a region involved in receptor binding selectivity^52^. This mutation may alter the function of the Ednrba protein, potentially affecting the dissociation of the endothelin ligands from the receptor^52–54^. Such changes could influence ENCC migration in cavefish. However, we found that *ednrba* is not expressed in migrating ENCCs as we did not observe *phox2bb* transcripts and *ednrba* transcripts in the same cells at 36 hpf and 48 hpf using colocalization analysis (BIOP version of JaCoP^55^ in ImageJ, Fig 3A-B). While we did not find evidence that *ednrba* is expressed in migrating ENCCs, we did observe *ednrba* transcripts at the distal end of the developing gut, suggesting it may nevertheless play a role in gut development (Fig 4B, E). Comparing Ednrbb, we found that the Pachón cavefish protein has one substitution at the N-terminus, 2 substitutions within the disordered region and G protein couples receptor family 1 domain compared to the surface fish protein (Table 1). Morpholino knock down of *ednrbb*, but not *ednrba*, reduces enteric neuron number in zebrafish^40^. Examining the expression of *ednrbb* may therefore help distinguish the roles of the endothelin receptor genes in *A. mexicanus* gut development.

**Figure 3.**
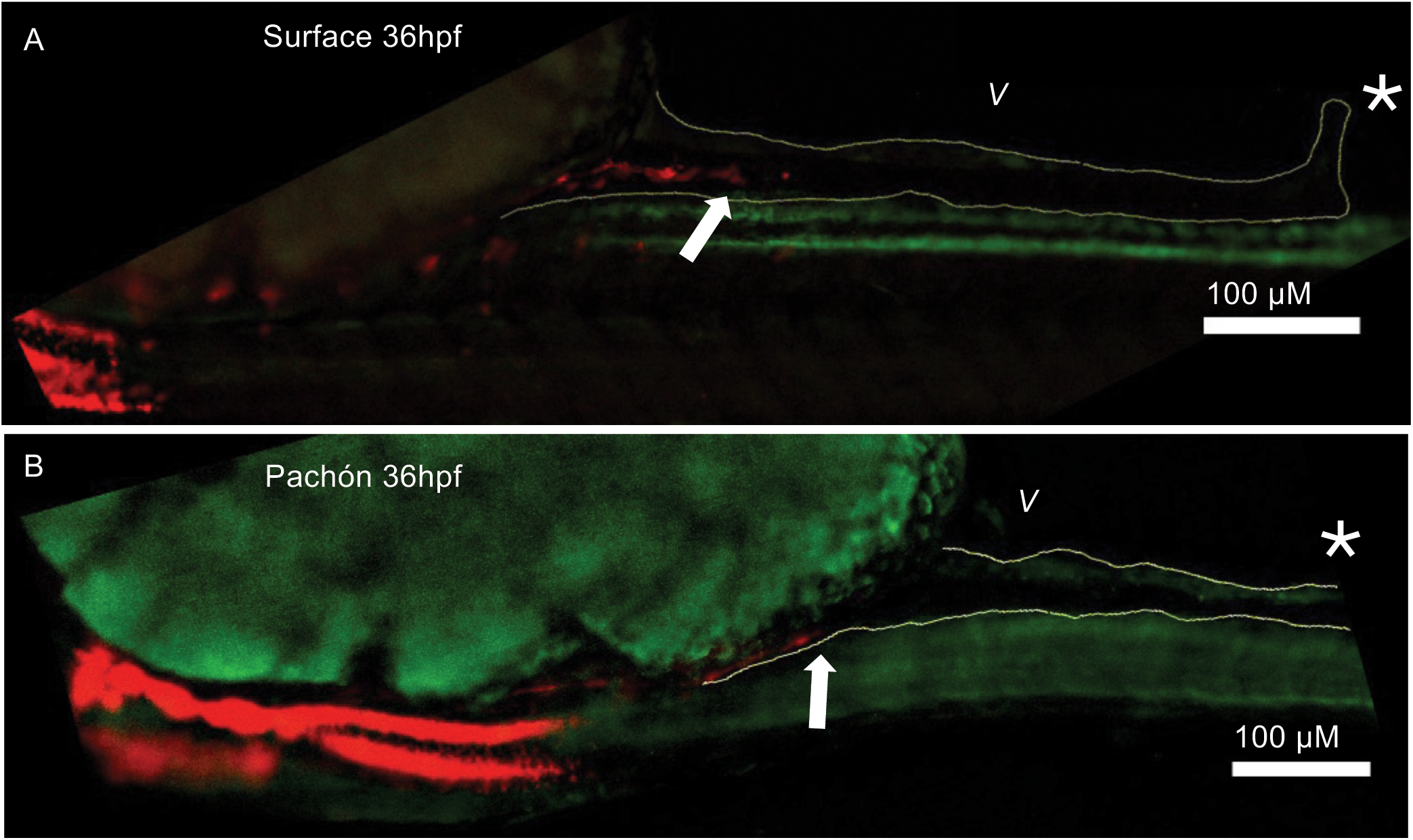
***Endothelin receptor-b* is not expressed in *phox2bb*-expressing enteric neural crest cells at 36 hpf in *Astyanax mexicanus.*** The expression of both *phox2bb* (red) and *ednrb* (green) can be observed in both Surface fish (A) and Pachón (B) larvae at 36 hpf; however, no colocalization was detected. Yellow outline – GI tract borders. White arrow – ENCCs front.. “*” Posterior lateral end of the larval fish gut. *“V”* Ventral side of the fish.

**Figure 4.**
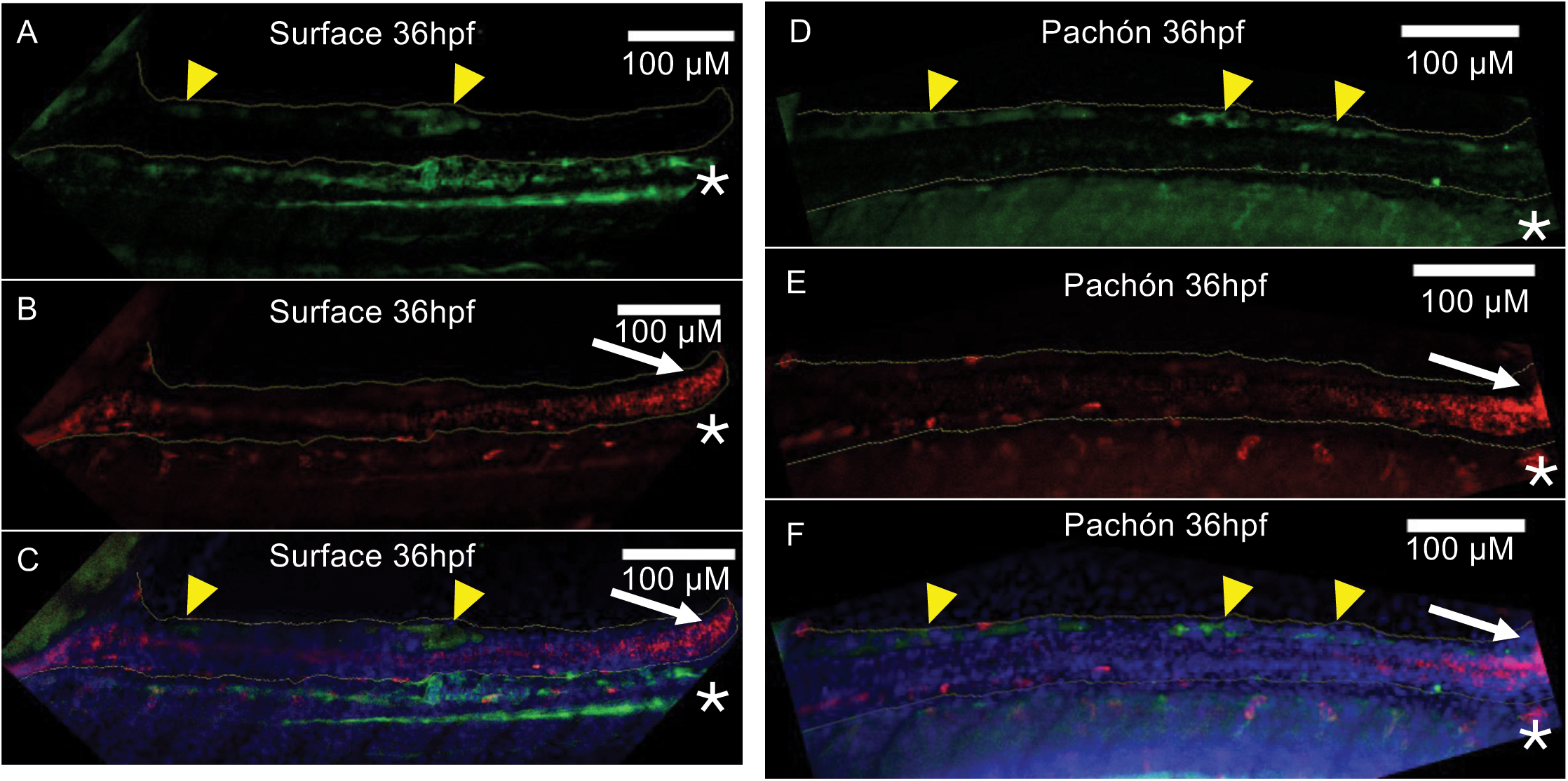
**Expression of endothelin-3 and endothelin receptor-b during enteric neural crest migration at 36 hpf in *A. mexicanus.*** *edn3* is expressed in a group of cells ventral to the developing gut (yellow arrow heads) posterior to presumptive migratory enteric neural crest cells at 36 hpf in both surface fish (A) and Pachón cavefish (D). *ednrb* is expressed in the distal gut end (white arrows) at 36 hpf in both surface fish (B) and Pachón cavefish (E). Overlay of signal from *edn3* and *ednrb* HCR probes for surface fish (C) and Pachón cavefish (F). Yellow outline – GI tract border. Blue – DAPI, Green – *edn3*, Red – *ednrb* “*” Posterior lateral end of the larval fish gut. Ventral side of the fish is toward the top of the images.

**Table 1.**
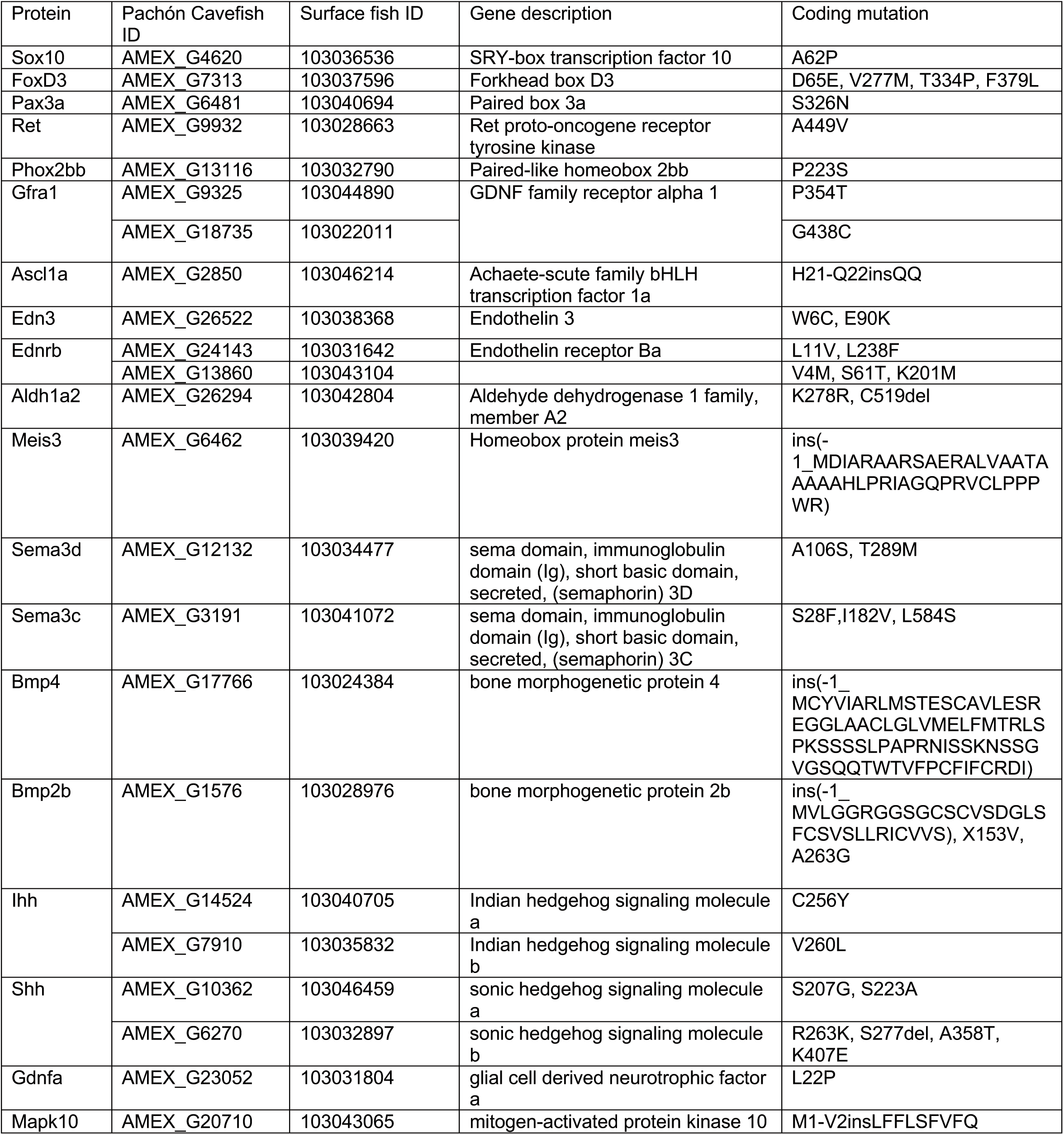
cavefish protein mutations in genes involved in ENS development.

Edn3, the Ednrb ligand, is secreted by the gut mesenchyme during ENCC migration and promotes ENCCs proliferation and migration while inhibiting differentiation^38,39,45^. *A. mexicanus* have two *edn3* genes, one on chromosome 13 (NCBI ID 103042214, AMEX_G16514) and one on chromosome 24 (NCBI ID 103038368, AMEX_G26522). The *edn3* gene on chromosome 13 is identical comparing surface fish and Pachón cavefish. In contrast, the *edn3* gene on chromosome 24 contains two substitutions in Pachón cavefish (W6C, E90K, Table 1). The E90K substitutions could influence proteolytic processing of the final peptide as it occurs near the cleavage site. We generated HCR probes that target the transcripts of the *edn3* gene located on chromosome 24. In 36hpf embryos, we observed cells expressing this gene in the gut posterior to migrating ENCCs, consistent with a role for *edn3* in promoting ENCC migration in *A. mexicanus* (Fig 4A, D).

Studying the role of *edn3* in ENCC migration utilizing the natural variation in *edn3* protein sequences in *A. mexicanus* could lead to a better understanding endothelin signaling in ENS development. Investigating the expression patterns of all Ednrb genes, analyzing the binding kinetics of the Ednrb with Edn3 variants, and conducting functional studies *in vivo* utilizing gene editing will clarify the role of Ednrb and its interaction with Edn3 in ENCC migration and differentiation in surface fish and Pachón cavefish.

### Cavefish display mutations in genes that regulate ENS development

To discover genetic changes that may result in more rapid ENCC migration and differentiation in Pachón cavefish compared to surface fish, we compared the predicted coding sequences of known regulators of ENS development (Table 1). We focused on key genes in the zebrafish ENS gene regulatory network (GRN), including *pax3a*, *foxd3*, *sox10*, *phox2bb*, and *ret*^31,32,35,56–59^. Pax3a and Foxd3 activate expression of *sox10* in migratory NCCs, which is necessary for expression of *phox2b* and *ret* in ENCCs^60,61^. Foxd3 also activates expression of the basic helix loop helix transcription factors *ascl1a* and *hand2* that coordinate differentiation of neuronal subtypes^62–66^. Genetic ablation of *pax3a*, *foxd3*, or *sox10* results in loss of ENCC colonization of the gut, while manipulation of more downstream GRN members like *phox2b* and *ret* reduces the final number of enteric neurons. We also compared the coding sequences of genes that work in combination with Ret including the intracellular kinase Mapk10, the secreted protein Sema3c/d, the chemoattractant Gdnfa, and the GDNF receptor Gfra1a/b^31,67–69^. In addition, we examined signaling molecules like retinoic acid (RA), sonic hedgehog (Shh), and bone morphogenetic protein (Bmp) that when mutated in zebrafish result in a range of ENS phenotypes, from total loss to reduced number of enteric neurons^70–72^. The results of the analysis are summarized in Table 1. Specific details on the mutations are described in the supplemental text, and a more extensive list of comparisons can be found in supplemental table 1.

Our comparative genomic analysis revealed mutations in cavefish proteins essential for ENS development, including amino acid substitutions and deletions in *pax3a, sox10, phox2bb, and foxd3,* as well as downstream targets like *ascl1a*. While these mutations often do not occur in predicted functional domains, they may still impact protein structure and function. We observed several changes in proteins related to the Ret signaling pathway. For example, Mapk10 showed differences in the number of splice isoforms comparing morphotypes, and Pachón Sema3c/d contained mutations within the Sema domain. Additionally, we identified variations in the chemoattractant Gdnfa and its receptor GFRA1, which may influence ENS cell migration and differentiation. Furthermore, mutations we observed in Aldh1a2, a critical enzyme in retinoic acid (RA) synthesis, suggest that RA signaling might be altered in cavefish, which could change the rate of ENCC migration and differentiation. We also observed coding changes within the Hint domains of sonic hedgehog genes (*shha* and *shhb*) that could influence ENCC proliferation and differentiation. In addition, we identified mutations in *bmp2b* and *bmp4*, key BMP signaling proteins, with insertions and substitutions that could affect neuron differentiation and ganglion formation. Our results provide a starting point for future functional studies and underscore the diverse genetic changes that may have contributed to the evolution of the ENS in cavefish, reflecting their adaptation to unique food sources in a subterranean environment.

## Conclusion

Using *Astyanax mexicanus* as an evolutionary developmental model, we established the developmental timeline for ENCC migration in surface fish and cavefish morphotypes. We observed that vagal neural crest cells begin migration at 24-27 hpf in both morphotypes, reaching the developing gut by 36 hpf. A notable difference emerges by 48 hpf: in surface fish, ENCCs have migrated only halfway through the developing gut, whereas in Pachón cavefish, they have already reached the hindgut, with differentiated enteric neurons present in the anterior gut. Additionally, *edn3* is expressed ahead of the migratory ENCCs in cells ventral to the gut tube in both morphotypes. However, no colocalization was found between *ednrba* and *phox2bb* in ENCCs at 36 hpf, indicating that the migratory ENCCs do not express *ednrba* in either morphotype. Finally, we identified mutations in candidate genes that may contribute to accelerated ENCC migration and differentiation. This study highlights how *A. mexicanus* can serve as a ‘natural mutant’ model for ENS development, offering insights into how the ENS evolves as vertebrates adapt to new environments and diets.

## Acknowledgments

We would like to acknowledge the laboratories of Johanna Kowalko (Lehigh University) and Clifford Tabin (Harvard Medical School) for providing fish embryos, Heather Woodson-Gammon (University of Nevada, Reno) and Brian Martineau (Harvard Medical School) for providing fish husbandry, and Megan Martik (Berkley) for HCR reagents and training. This research was supported by a grant from the National Institutes of Health (P30 GM145646).

## Declaration of generative AI and AI-assisted technologies in the writing process

During the preparation of this work the author(s) used ChatGPT to improve readability of the manuscript. After using this tool/service, the author(s) reviewed and edited the content as needed and take(s) full responsibility for the content of the publication.

---

**Supplementary Table 1.**
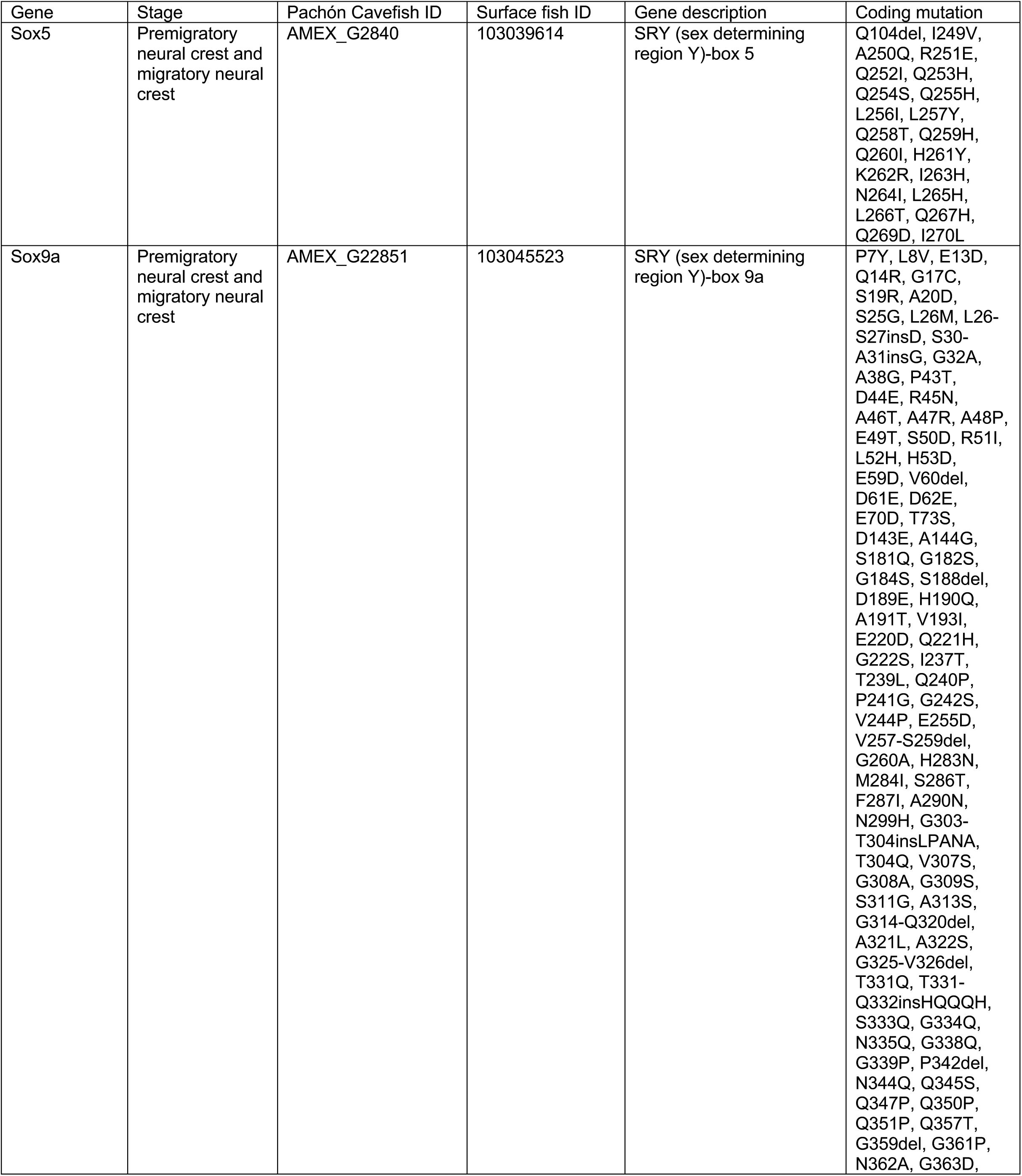
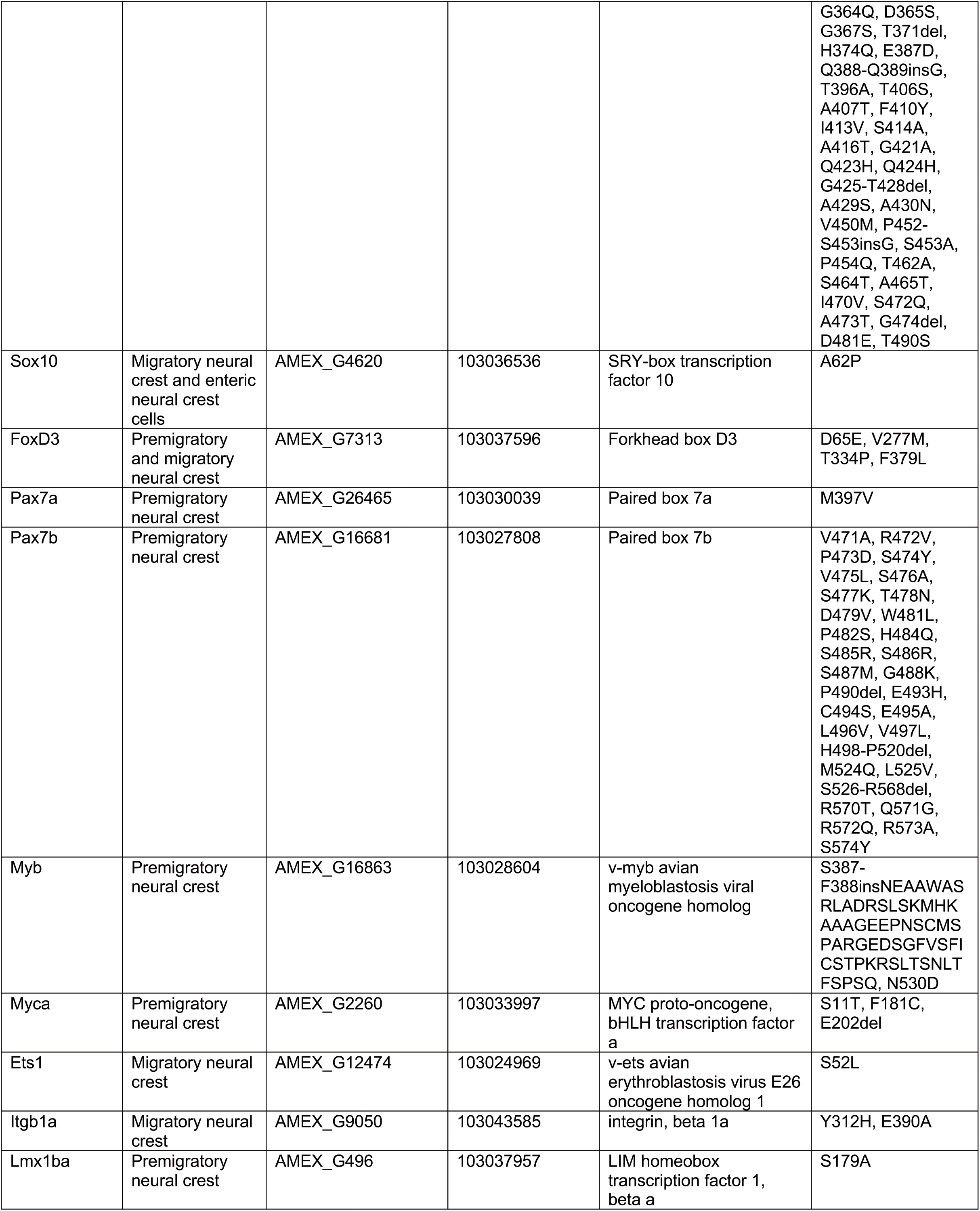
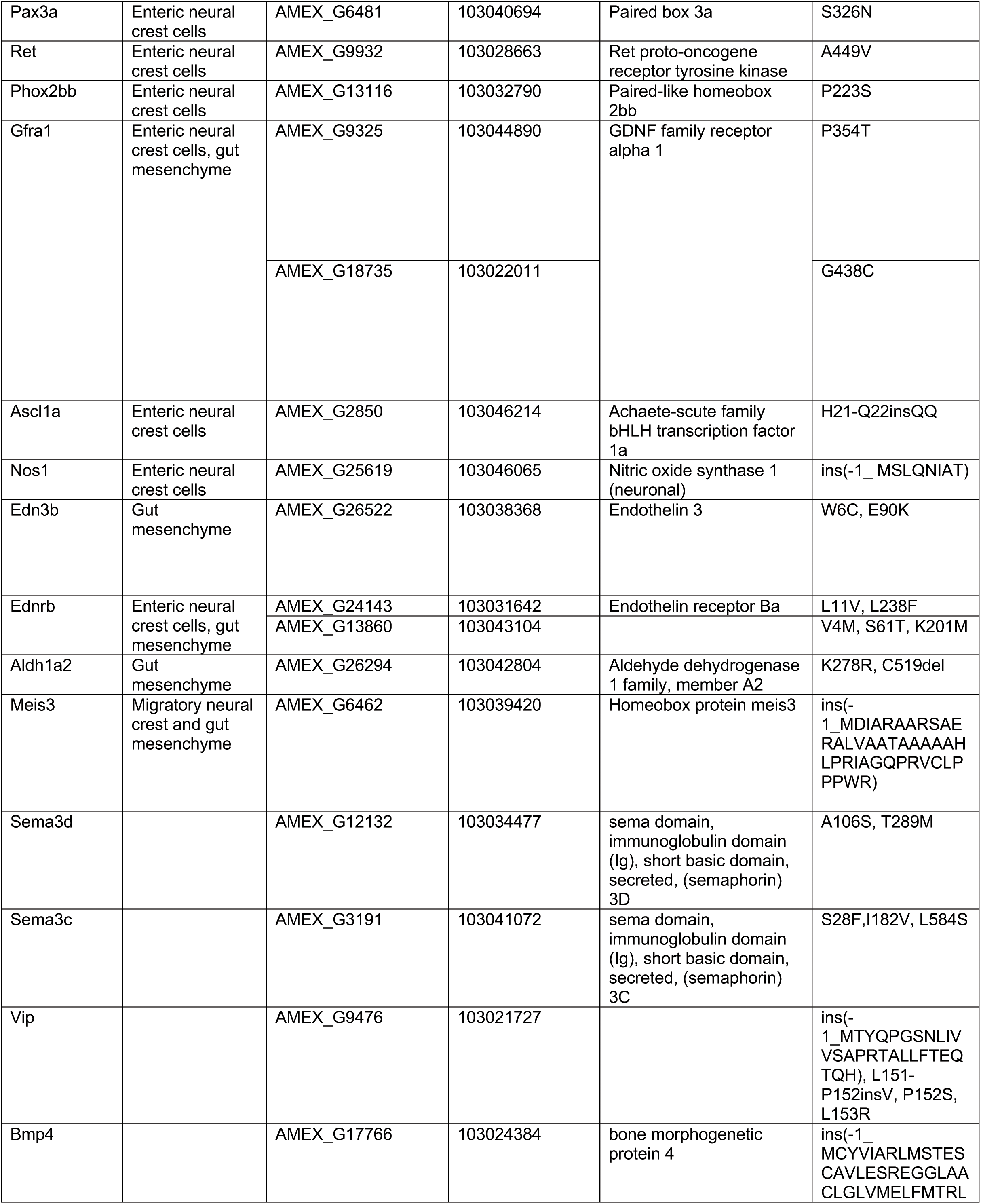
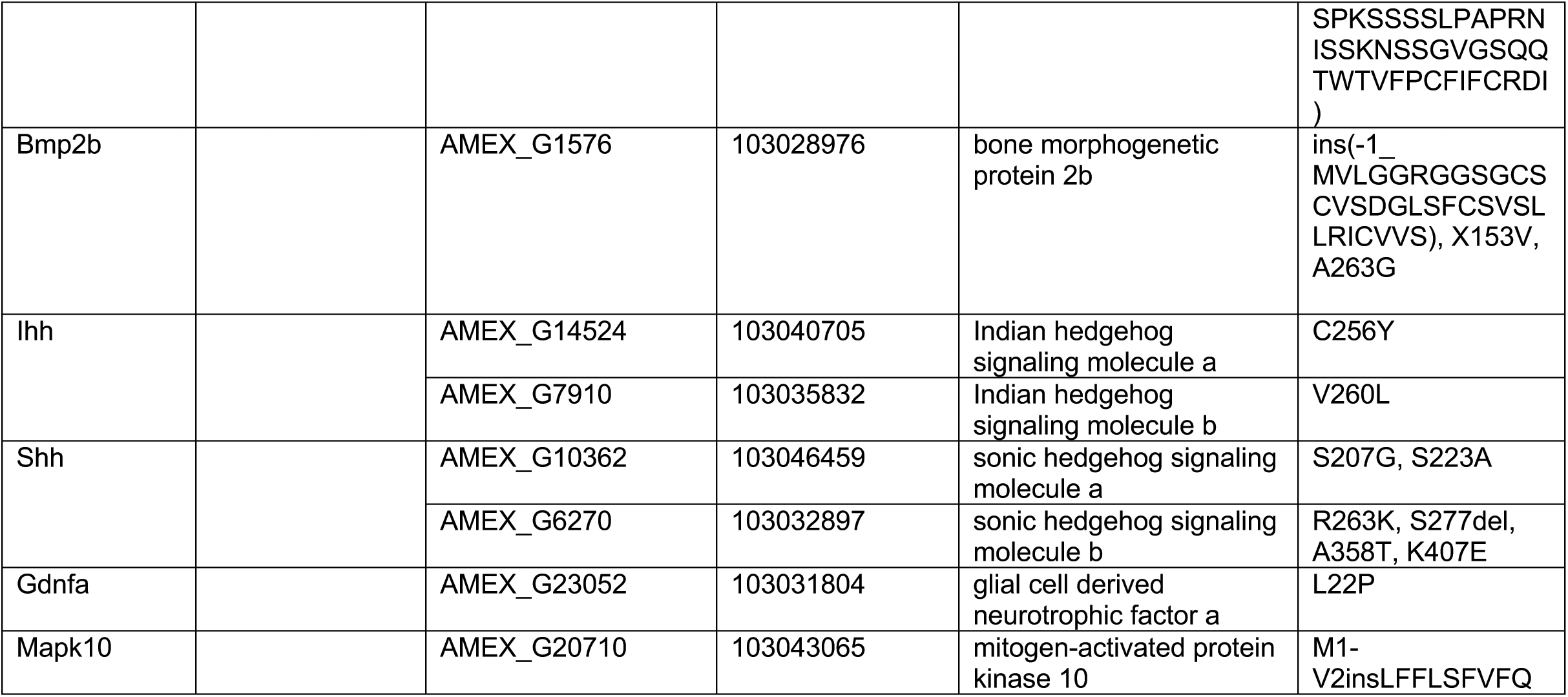
cavefish protein mutations in genes involved in ENS development.

## Supplemental text

To discover genetic changes that may result in more rapid ENCC migration and differentiation in Pachón cavefish compared to surface fish, we compared the predicted coding sequences of known regulators of ENS development (Table 1). We discovered predicted coding mutations in the proteins important for initiating the ENS GRN and coordinating later stages of ENS development in Pachón cavefish. For example, Pax3a contains one amino acid substitution (S326N) occurring in the disordered region of the protein. Sox10, Ret, and Phox2bb proteins each contain one amino acid substitution (A62P, A449V, P223S respectively) that do not occur in predicted functional domains but within the polar residue compositional bias regions. Foxd3 contains four amino acid substitutions (D65E, V277M, T334P, F379L) including one substitution (D65E) within the predicted fork head domain. The downstream transcriptional targets of Foxd2, *ascl1a*, also possess mutations in Pachón cavefish compared to surface fish. Ascl1a exhibits a 10 amino acid deletion and 10 substitutions. These variants do not occur in predicted functional domains of the proteins, however.

While Ret only contains one amino acid substitution in Pachón cavefish, other players in the Ret network contain numerous coding variants. Mapk10 is predicted to have five splice isoforms in surface fish and one in Pachón cavefish. Comparing the longest surface splice isoform with Pachón protein (NCBI ID 103043065, AMEX_G20710), there is an insertion at the N-terminus of the Pachón Mapk10 protein sequence. Pachón cavefish Sema3c has three amino acid substitutions including one (I182V) located within the Sema domain which may impact the function of the protein. There are two predicted splice isoforms for Sema3d in surface fish while Pachón cavefish genome has only one isoform. We compared the longest splice isoforms from surface and Pachón cavefish (NCBI ID 103034477, AMEX_G12132) and found that the predicted peptide in Pachón contains two single amino acid substitutions (A106S, T289M) that occur in the Sema domain (AA64-553). Comparing the second surface fish sema3d isoform with Pachón protein sequence, in addition to the above mentioned two amino acid substitutions, Pachon protein has an insertion (K536-Q537insQ) within the same Sema domain.

The Pachón protein sequence of the chemoattractant, Gdnfa, has one amino acid substitution (L22P) at the N-terminus compared to the surface fish protein. GFRɑ1 activates GDNF-bound Ret^73^. Morpholino knockdown of gfra1a and gfra1b in zebrafish leads to reduced enteric neurons^31^. Zebrafish *grfra1a* is orthologous to surface fish *gfra1a* (NCBI ID: 103044890) while gfra1b is orthologous to *gfra1b* (NCBI ID: 103022011). There are two annotated *gfra1* genes in the Pachón cavefish genome: *GFRA1* (NCBI ID: AMEX_G9325), and *GFRA1-like* (NCBI ID: AMEX_G18735). Pachón GFRA1 protein is orthologous to surface fish gfra1a and contains a single amino acid substitution (P354T) in the predicted GDNF/GAS1 domain of the protein. Moreover, Pachón GFRA1-like protein is orthologous to surface gfra1b protein. Pachón GFRA1-like protein has one amino acid substitution at the C-terminus (G438C) compared to the surface fish gfra1b protein sequence.

Retinoic acid (RA) is crucial for the migration and survival of ENCCs during gut development^70,74,75^. RA acts by regulating key genes such as *meis3* and *ret* receptor, ensuring that ENCCs migrate properly and colonize the gut completely^70^. The retinaldehyde dehydrogenase enzyme Aldh1a2 catalyzes the formation of RA (reviewed in^76–78^). Comparing the predicted protein sequences between surface fish and Pachón cavefish, Pachón protein contains one deletion (C519del) at the C-terminus of the protein and one amino acid substitution (L278R) that occur within the Aldehyde dehydrogenase protein domain (AA47-510) suggesting they may impact enzyme function and therefore RA production. In addition, Pachón cavefish have yellow yolk compared to surface fish indicating an accumulation of the RA precursor molecules, carotenoids, suggesting other steps of RA synthesis may be altered in cavefish^19^. Studying how evolution of retinoic acid signaling has impacted cavefish neural crest development represents a promising avenue for future discoveries.

The co-transcriptional regulator Meis3 may connect RA and Shh signaling; reduced production of RA is associated with lower meis3 expression, and loss of Meis3 function prevents intestinal *shha* expression^70,79^. Zebrafish meis3 is orthologous to surface fish (NCBI ID: 103039420) and Pachón cavefish (NCBI ID: AMEX_G6462) meis3. Pachón cavefish meis3 protein has a 41 amino acids insertion at the N-terminal of the protein sequence. However, the homeobox domain and disordered regions are well conserved between the two morphotypes, likely preserving the protein’s function.

Both sonic hedgehog (shh) and Indian hedgehog (ihh) are involved in development of the ENS in vertebrates^72,80–82^. Endoderm-derived Shh is essential for the radial patterning of the ENS^80,82^, promoting the proliferation of ENCCs while inhibiting their neuronal differentiation and restricting their GDNF-induced migration^81^. In the absence of ihh, NCCs can migrate and differentiate but fail to survive or proliferate, underscoring ihh’s critical role in ENS development^80^. In zebrafish, endoderm-derived shha drive both initial migration of ENCCs from the vagal region to the developing gut and their subsequent migration along it^72^. In addition, shha functions as a mitogen, promoting the proliferation of ENS precursors, and is essential for the expression of gdnf in the intestinal mesenchyme^72^. In surface fish, shh expression follows restricted pattern, particularly along the ventral midline of the neural plate. In cavefish, however, shh expression is particularly expanded and elevated throughout development^83–86^. However, the expression patterns and differences of gut endoderm-derived shh in *Astyanax mexicanus*, as well as the impact of expanded shh expression along ventral midline of the neural plate on neural crest and ENS development, remain unknown.

Both Surface fish and Pachón cavefish genome possess distinct two shh isoforms; shha (NCBI ID: 103046459, AMEX_G10362) and shhb (NCBI ID: 103032897, AMEX_G6270) which are orthologous to the zebrafish shha (ENSDARG00000068567) and shhb (ENSDARG00000038867) isoforms. The shha protein in Pachón cavefish has two amino acid substitutions (S207G, S223A) within the Hint domain, compared to the surface fish shha protein. The Pachón cavefish shhb protein exhibits a single substitution (R263K) and a deletion (S277del) within the Hint domain and an amino acid substitution (K407E) at the C-terminus on the protein relative to the surface shhb protein. These multiple alterations in the functional domains of both shha and shhb in Pachón cavefish may affect the expression and function of the proteins during ENS development. We currently lack information on the expression patterns and differences of gut endoderm-derived *ihh* in *Astyanax mexicanus*. In zebrafish, *ihha* and *ihhb* are orthologous to *A. mexicanus* surface fish *ihha* and Pachón cavefish IHHB (NCBI ID: 103040705, AMEX_G14524) and surface fish *ihhb* and Pachón cavefish IHHB-like (NCBI ID: 103035832, AMEX_G7910), respectively.

In Pachón cavefish, the ihha protein contains a single amino acid substitution (C256Y) within the Hint domain. The ihhb protein in Pachón cavefish features a single substitution (V206L) within the Hint functional domain, which may alter the domain’s structure and lead to functional differences.

Bone morphogenetic protein (BMP) signaling is activated in the ENS cells during various stages of gut development^87,88^. Misexpression of BMP-4 led to abnormalities in ENS development, including altered ganglion formation and positioning, indicating that precise regulation of BMP signaling is necessary for normal ENS patterning and neuron differentiation^87^. BMP-2 and BMP-4 were found to limit the overall number of enteric neurons. High concentrations of these proteins led to a reduction in the number of neurons, partly due to increased apoptosis and decreased proliferation^71^. While the functional role of BMP signaling in craniofacial^89^ and teeth/jaw development^90^ has been well studied in *A. mexicanus*, the expression and functional roles of BMP secreted by the gut mesoderm and endoderm in relation to ENCCs and ENS development remain largely unexplored. Zebrafish has two isoforms of bmp2; bmp2a and bmp2b which are orthologs for A. mexicanus bmp2a (NCBI ID: 103028976, AMEX_G17740) and bmp2b (NCBI ID: 103028976, AMEX_G1576). bmp2b is a critical component of the zebrafish gene regulatory network^37^. The protein sequences of both surface fish and Pachón cavefish bmp2a are identical. In Pachón cavefish, the bmp2b protein has two single substitutions before the disordered region and large insertion of 32 amino acids before the N terminus of the surface fish bmp2b protein. Compared to the surface bmp4 protein, the Pachón cavefish bmp4 protein has a 79-amino acid insertion at the N-terminus making it larger than the surface protein. In addition, we observed coding changes in additional genes that are involved in ENS development as summarized in supplementary table 1.

